# A model for adult organ resizing demonstrates stem cell scaling through a tunable commitment rate

**DOI:** 10.1101/137638

**Authors:** XinXin Du, Lucy Erin O’Brien, Ingmar Riedel-Kruse

## Abstract

Many adult organs grow or shrink to accommodate different physiological demands. Often, as total cell number changes, stem cell number changes proportionally in a phenomenon called ‘stem cell scaling’. The cellular behaviors that give rise to scaling are unknown. Here we study two complementary theoretical models of the adult Drosophila midgut, a stem cell-based organ with known resizing dynamics. First, we derive a differential equations model of midgut resizing and show that the in vivo kinetics of growth can be recapitulated if the rate of fate commitment depends on the tissue’s stem cell proportion. Second, we develop a twodimensional simulation of the midgut and find that proportion-dependent commitment rate and stem cell scaling can arise phenomenologically from the stem cells’ exploration of physical tissue space during its lifetime. Together, these models provide a biophysical understanding of how stem cell scaling is maintained during organ growth and shrinkage.

## Introduction

Mature organs contain both differentiated cells, which execute physiological function, and stem cells, which generate new differentiated cells. In organ homeostasis, stem cells divide to replace differentiated cells that are lost, and numbers of stem and differentiated cells are constant. Increased functional demand can induce adaptive growth, a transient, non-homeostatic state in which stem cells divide to generate excess differentiated cells [1–4]. Similarly, decreased demand leads to adaptive shrinkage, in which differentiated cells are reduced in part because stem cells cease to divide [5, 6]. Adaptive resizing enables mature organs to maintain physiological fitness in the face of changing environmental conditions [1, 7–9].

Intriguingly, many organs exhibit altered numbers of stem cells in response to major physiological adaptation or resizing; examples of these include altered numbers of satellite stem cells in muscles after exercise or induced hypertrophy [10–12], altered numbers of mammary gland stem cells during pregnancy [13, 14], and altered numbers of intestinal stem cells after feeding [15]. In particular, authors in [15] found that stem cells scale with the size of the organ, that is, stem cells adjust their numbers during resizing to remain a similar proportion of total cells in the organ. Because of scaling, the cellular ‘replacement burden’ of an individual stem cell stays constant irrespective of organ size. Physiologically, the constant replacement burden may be advantageous since it allows the organ to respond exponentially quickly (at least initially) to environmental changes. This is because the rate of change of the size of the system would typically be proportional to the number of stem cells and therefore proportional to the size of the system, leading to exponential response. In the adult Drosophila midgut, a simple epithelial organ functionally equivalent to the vertebrate small intestine, this scaling behavior is extraordinarily precise; a four-fold increase in differentiated cells, induced by increased dietary load, is matched by a four-fold increase in stem cells [15]. Importantly, for stem cell scaling to occur, there must be population-wide coordination between symmetric and asymmetric fate outcomes after cell division [15–17]. While we know that at the individual cell level, fate outcomes are determined through Delta-Notch signaling, we do not know what mechanisms coordinate stem cell scaling at the population level.

Some prior models of the midgut [16, 18] and of other self-renewing organs [18–27] have considered homeostasis without adaptive resizing. Some of these as well as other models have considered embryonic development [17, 18, 28, 29] or cancer [19, 24, 27, 30–32], two growth states that do not exhibit stem cell scaling. To shed light on scaling mechanisms, we develop a set of non-spatial differential equations as well as a two-dimensional simulation of cell dynamics in the Drosophila midgut. Here we find that the physiological kinetics of stem cell scaling during midgut adaptive growth can be recapitulated by a set of ordinary differential equations. The ability of these equations to recapitulate physiological kinetics depends strongly on the inclusion of feedback. Specifically, physiological dynamics of cell populations are captured if the rate at which new cells commit to differentiation depends on the existing proportion of stem cells. Next, we develop a two-dimensional simulation of the midgut and show that this tunable commitment rate can be explained by the concept of a ‘stem cell territory’ – the physical space that a stem cell explores during its lifetime. We show that territory size is determined by cell-cell adhesion, stochastic motility, and Delta-Notch signaling. We find that stem cell scaling requires a threshold territory size and that systems within this regime fit our differential equations.

## Non-spatial Model

### Description of non-spatial model

We propose a mathematical description of the midgut that considers the midgut’s three major cell types: stem cells, enteroblasts, and enterocytes, each of which exhibits a distinct cellular behavior (Figure 1). Stem cells, *s*, are the only cells in the midgut that typically divide [33]. In our model, all stem cell divisions generate two daughters that are equipotent stem cells [16]; this division rate is denoted *a*. Some stem cells become enteroblasts, *u*. Enteroblasts are post-mitotic and committed to differentiate into enterocytes, but still lack the morphological features of differentiation. The rate that stem cells commit to terminal fate is denoted *b*. Enterocytes, *U*, are fully mature epithelial cells that comprise most of the cells in the midgut epithelium. Enteroblasts differentiate into enterocytes at a rate *λ*. Enterocytes die, or are otherwise lost, at a rate Λ.

**Figure 1:**
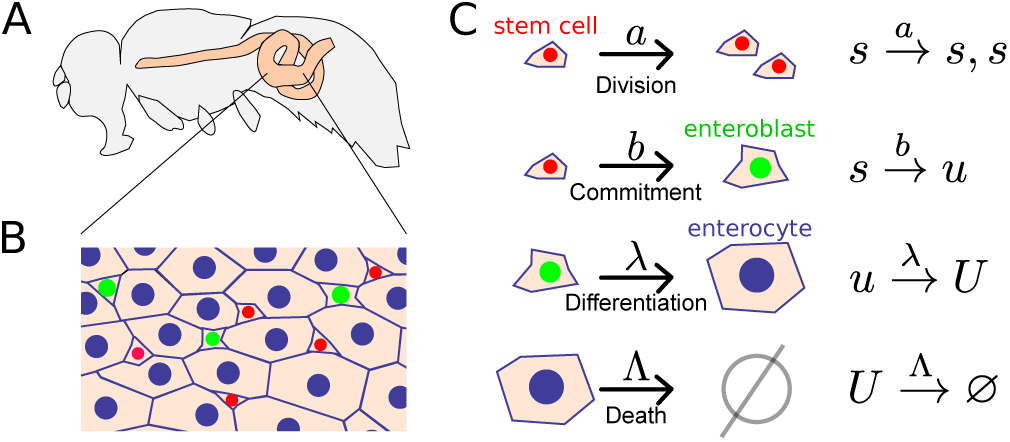
Stem cells in the Drosophila midgut regulate growth and homeostasis. (A) Schematic of the Drosophila midgut (yellow). (B) Schematic of midgut epithelium with stem cells (red) and enteroblasts (green) situated among enterocytes (blue outline). (C) Cellular processes taken into account by the model in Equation 1.

These cellular relationships lead to a simple mathematical description for numbers of stem cells, enteroblasts, and enterocytes as functions of time *s*(*t*), *u*(*t*), and *U* (*t*) with the form:

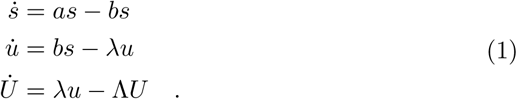

For the system to have a non-zero, finite steady state, either rates of division and commitment must always be equal (*b* = *a*), which would imply highly precise regulation of these two processes, or else some rates in Figure 1 and Equation 1 must depend on cell numbers. We assume the latter case since highly precise regulation is unlikely in a noisy tissue system.

To account for nutrient-driven adaptive growth and shrinkage [15], we propose that cell numbers depend on ingested nutrients, and that the system contains feedback. We denote the total energy of ingested nutrients as *E*_in_ and define the ‘energy density’ *E*_*d*_ as:

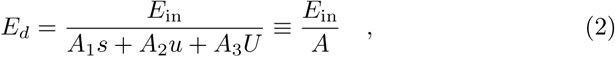

in which *A ≡ A*_1_*s* + *A*_2_*u* + *A*_3_*U* is the total tissue consumption of *E*_in_, and *A*_1_, *A*_2_, and *A*_3_ denote the cell-type specific consumption per cell.

Physiologically, we assume that cells most likely can access only the local value of nutrients, namely the energy density. Therefore, we designate rates of cell division *a* and cell loss Λ to depend on *E*_*d*_. In fact, *In vivo* analysis of the midgut has shown that high levels of nutrients promote divisions and suppress death, whereas low levels of nutrients has the opposite effect, as reflected by levels of insulin signaling [15]. Accounting for these observations, denoting *a*_*m*_ and Λ_*m*_ as maximal rates of division and death, respectively, non-dimensionalizing *E*_*d*_, and choosing Hill functions to represent generic sigmoidal functions, we arrive at nutrient-dependent expressions for division and death rates:

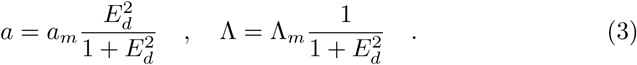

Note that although we chose second order Hill functions to represent the sigmoidal dependence of cell division and loss rates on nutrient density, other Hill functions do not significantly alter the results, as detailed in the Supplement. Model equations for *s*, *u*, and *U* taking into account nutrient-dependent division and death rates are therefore:

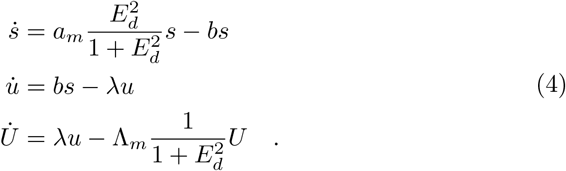

Importantly, the steady states of Equation 4 satisfy scaling because {*E*_in_, *s, u, U* } → {*αE*_in_, *αs, αu, αU*}leaves the steady state equations for Equation 4 invariant.

### Constant versus stem cell proportion-dependent commitment rate: compatibility with empirical measurements

We next sought to compare the solutions of Equation 4 to the known dynamics of the midgut *in vivo*. Solving Equation 4 under steady state conditions, we find for steady state stem cell number ratios *s*_0_*/u*_0_, *s*_0_*/U*_0_, and *s*_0_*/*(*s*_0_ + *u*_0_ + *U*_0_):

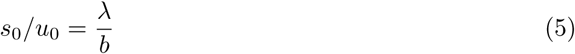

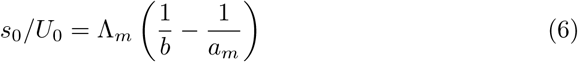

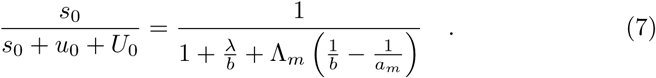

Full steady state solutions to Equation 4 are in the Supplement. Note that in this general form, we assume that *b* may have functional dependence on *s*_0_, *u*_0_, or *U*_0_ and that b cannot surpass *a*_*m*_ in value.

Important to the discussion of empirical measurements in the midgut is the definition of symmetric and asymmetric fate outcomes. Since, as a first approximation, stem cells are the only cells in the midgut capable of dividing [33], we designate fate outcomes as *symmetric-stem* if both daughter cells divide before either daughter becomes a committed enteroblast. Following similar logic, we designate fates as *symmetric-terminal* if both daughters become committed enteroblasts before either divides, and as *asymmetric* if one daughter becomes a committed enteroblast while the other divides [15].

Since we model division and commitment as Poisson processes (uncorrelated) with rates *a* and *b*, the probability that a given stem cell undergoes division before commitment is *a/*(*a*+*b*). The frequency of symmetric-stem fate outcomes is therefore

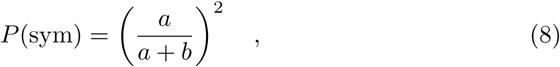

where *a* and *b* can have any functional dependence. Note that, in steady state, similar to the results in [18], we have *P* (sym) = 1*/*4 because *a* = *b* in steady state. The maximum symmetric-stem division rate, often achieved during growth, is *P*_max_(sym) = (*a*_*m*_*/*(*a*_*m*_ + *b*))^2^.

#### Constant rate of commitment is incompatible with empirical measurements

Other models of tissue homeostasis have assumed that commitment rate is constant [16, 22]. We thus asked whether a constant commitment rate is compatible with growth. Solving Equation 4 for *b* = *B*_0_ = constant and a 4x increase in food, we find that the key features of midgut growth *in vivo* are indeed recapitulated: cell numbers increase, the frequency of symmetric-stem fates increases transiently, and stem cell number scales (Supplement). Here, the value of 4x food increase was chosen to generate a 4x increase in cell numbers so as to best compare to the experimentally measured 4x increase in cell numbers when the gut undergoes feeding [15]. As noted, the cell number ratios *s*_0_*/U*_0_ and *s*_0_*/u*_0_ are independent of food input or absolute cell number, so stem cell scaling arises naturally. Thus in principle, a constant commitment rate is compatible with resizing.

However, the validity of a constant commitment rate model breaks down when we compare the theoretical parameter space with the biologically observed parameter space. Although commitment rates have not been experimentally measured, it is known that in steady state, *s*_0_*/u*_0_ *≈* 1 and *s*_0_*/U*_0_ *≈* 0.2 during midgut homeostasis [15, 16]; specifically, the numbers of stem cells, enteroblasts and enterocytes are around 700 *±*200, 500*±* 200, and 2800 *±* 800, respectively [15]. Using these values in Equations 5 and 6, and replacing commitment rate *b* with the constant *B*_0_, we obtain:

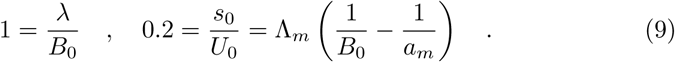

Using these relationships to deduce *λ* and *B*_0_ from *a*_*m*_ and Λ_*m*_, we solve Equation 4 for various values of *a*_*m*_ and Λ_*m*_. To focus our parameter range, we apply three criteria. First, homeostatic stem cell division rates *a*_0_ vary from 0.5 times/day to 4 times/day [15, 16]. Given that division and commitment must be equal at homeostasis (*a*_0_ = *B*_0_), this biologically relevant range of *a*_0_ is obtained from *a*_*m*_ and Λ_*m*_ using Equation 9 (Figure 2A, blue shading). Second, the maximum reported rate of symmetric-stem fates under physiological conditions is 0.7, which is transiently observed during growth [15]. We thus set *P*_max_(sym) = (*a*_*m*_*/*(*a*_*m*_ + *b*))^2^ *≈* 0.7 (with *b* = *B*_0_) and included a margin of error such that 0.5*≤ P*_max_(sym) *≤*0.9 (Figure 2A, red shading), where the margin of error is estimated from data in [15] (see Supplement). Third, the time required for the growth phase to reach completion is approximately 3.5 days [15], after which homeostasis is re-established at the organ’s new, larger size. Therefore, only values of *a*_*m*_ and Λ_*m*_ for which solutions to Equation 4 approach steady state by *t* = 3.5 days give rise to steady states with *in vivo* kinetics (Figure 2A, white region). Conditions used to determine whether Equation 4 has reached approximate equilibrium are in the Supplement.

**Figure 2:**
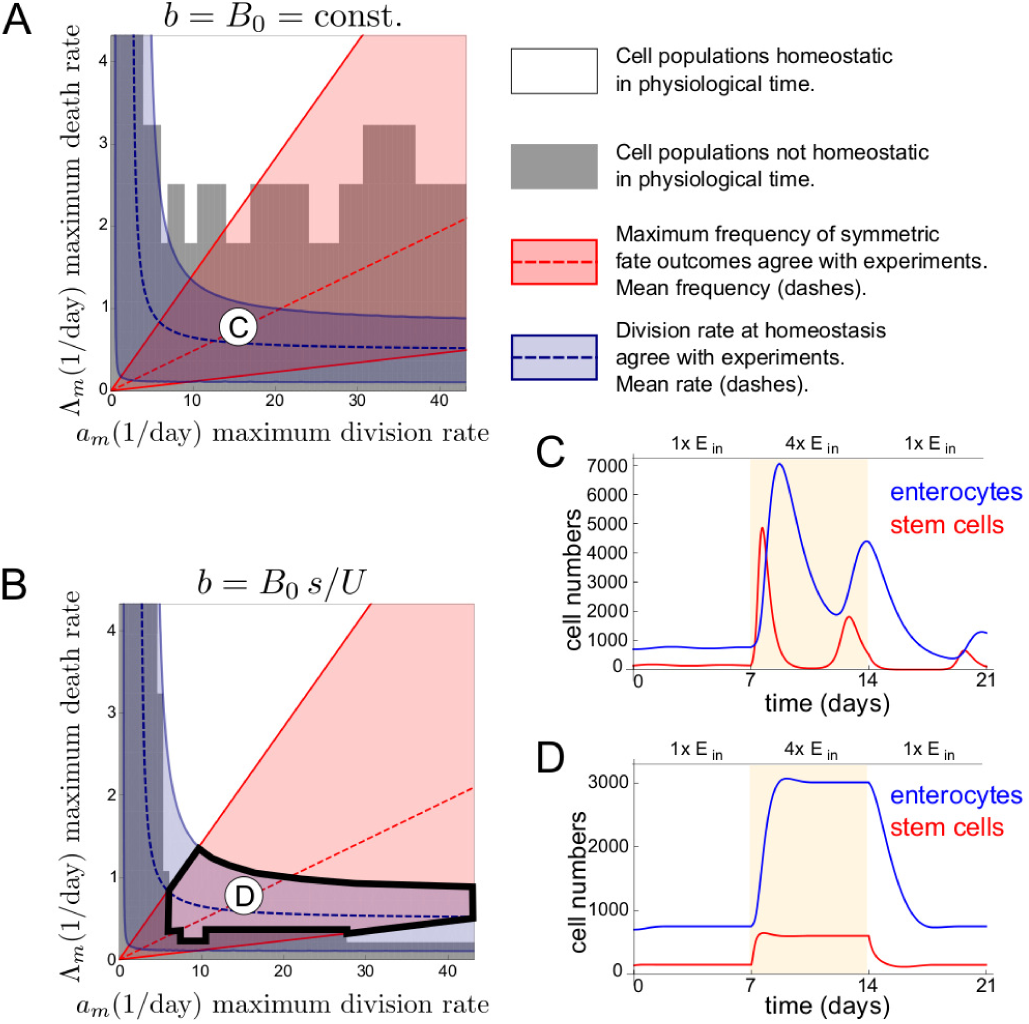
Stem cell proportion-dependent commitment rate *b* = *B*_0_*s/U* model satisfies experimental observations whereas constant commitment rate *b* = *B*_0_ model does not. (A,B) For *b* = *B*_0_ (A) and *b* = *B*_0_*s/U* (B), regions of parameter space (*a*_*m*_, Λ_*m*_) that satisfy: homeostatic division rate 0.5 /day *≤ a*_0_ *≤* 4.0 /day (blue), maximum frequency of symmetric-stem fate outcomes 0.5 *< P*_max_(sym) *<* 0.9 (red), and homeostasis within *t* = 3.5 days (white). Averages *a*_0_ = 2.5 /day and *P*_max_(sym) = 0.7 indicated by dashed lines. Parameter values satisfying all experimental measurements do not exist for *b* = *B*_0_ model (A), while they exist for *b* = *B*_0_*s/U* model (B) (solid black outline). Circled letters indicate parameters corresponding to solutions in panels (C-D). (C) Solutions of *b* = *B*_0_ model are highly oscillatory in parameter regimes that satisfy experimental values of division rate and maximum frequency of symmetric-stem fate outcomes. (D) Solutions of *b* = *B*_0_*s/U* model respond quickly to food changes without large oscillations in parameter regimes that satisfy all experimental measurements.

Applying these criteria, we observe that when commitment rate is constant (*b* = *B*_0_), no regions of the theoretical parameter space satisfy all three biological measurements; i.e., there is no region in which blue, red, and white shading all overlap (Figure 2A). Specifically, solutions to Equation 4 with constant *b* = *B*_0_ show strong behaviors of ringing (see Figure 2C and Supplement) which prevents solutions from approaching steady state within physiological time. This ringing in solutions approaching a new steady state determined by *E*_in_ is present for all fold-changes of *E*_in_. Thus, although constant commitment rate is theoretically compatible with midgut adaptive growth, it is incompatible when biologically relevant measurements are applied.

#### A rate of commitment that depends on stem cell proportion is compatible with scaling and empirical measurements

Since the model with constant commitment rate does not fit experimental measurements, we examine a second possibility, in which commitment rate varies depending on how many stem cells and enterocytes are present in the tissue. Such a relationship can give rise to more realistic dynamics by allowing cell numbers to feed back into the commitment rate. In the midgut, commitment to enteroblast fate is known to occur through Delta-Notch signaling between stem cells [33–36]. This biological phenomenology suggests that it is reasonable to model the signaling frequency, and consequently commitment rate, such that it depends on the number ratio of stem cells in the tissue *s/U*. We therefore explore the effects of this additional feedback to our model in Equation 4.

We find that a commitment rate proportional to stem cell-enterocyte ratio recapitulates midgut resizing within timescales that are biologically relevant. Specifically, we modify three elements: Equation 4 such that *b* = *B*_0_*s/U*, Equation 9 such that *B*_0_ is replaced by *B*_0_*s*_0_*/U*_0_, and the *P*_max_ calculation such that *b* = *B*_0_*s*_0_*/U*_0_. Incorporating these modifications, we solve Equation 4 for various values of *a*_*m*_ and Λ_*m*_. The resulting values define a theoretical parameter space that is compatible with the known biological measurements; i.e., the region in which blue, red, and white shading overlap (Figure 2B). In addition, stem cell scaling is maintained due to the form of *b* = *B*_0_*s/U*. Importantly, the large oscillations and ringing behaviors in the *b* = *B*_0_ model (Figure 2C) that delays the approach to steady state are suppressed in the *b* = *B*_0_*s/U* model (Figure 2D). This result is confirmed by linear stability analysis, detailed in the Supplement. Moreover, the parameter region in which oscillations are significant is smaller in the *b* = *B*_0_*s/U* compared to the *b* = *B*_0_ model, also detailed in the Supplement. We conclude that the known kinetics of midgut resizing can be accounted for by introducing an additional feedback term to Equation 4, in particular, one that tunes the rate of enteroblast commitment to the existing proportion of stem cells.

Therefore, from these results, we propose that our model equations be modified as:

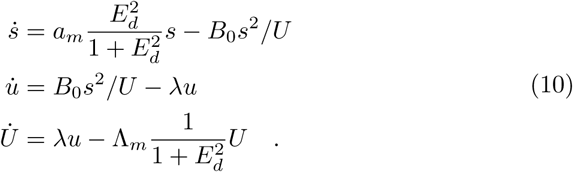

Equation 10 captures stem cell scaling and *in vivo* resizing dynamics of the midgut. The steady state solutions to Equation 10 are presented in the Supplement. We note that the feedback through stem cell number ratio *s/U* does not uniquely allow the resizing model in Equation 4 to capture realistic dynamics in the midgut; however, proportionality of commitment rate to *s/U* is one of the simplest forms for feedback that both represents biological phenomenology and captures realist dynamics. Alternative models for feedback are suggested and explored in the Supplement.

## 2-dimensional model

### 2-dimensional model description

Although our non-spatial model implies that a tunable commitment rate *b* is needed for midgut resizing that fits experimental measurements, it does not provide insight into the cellular mechanisms that underlie this tunability. To explore these potential mechanisms, we develop a two-dimensional model of the midgut that builds upon our non-spatial model. Many mathematical approaches have been used to model two-dimensional epithelia including vertex models [37–40], cellular automata [41–43], cellular Potts models [44, 45], and cell-centered models [46–48]. Our 2D model is a cell-centered model based on overlapping spheres [49–51], where we represent cells as 2D-spheres (disks) that are specified by the positions of their centers and that interact physically via forces such as cell-cell adhesion, volume exclusion, and stochastic motion (Figure 3A-B). We chose a cell-centered model, among others, because it describes the midgut system at the length scale of cells, which is the length scale of the phenomenology in which we are interested. As shown, *in vivo* [35, 52], contact-mediated, Delta-Notch signaling between individual stem cells serves to define commitment to terminal fate (Figure 3C). Thus, in the two-dimensional model, commitment rate becomes a property that emerges from the fate decisions of individual enteroblasts, in contrast to the non-spatial model, in which commitment rate is explicitly defined with respect to the tissue-wide ratio of stem cells to enterocytes.

**Figure 3:**
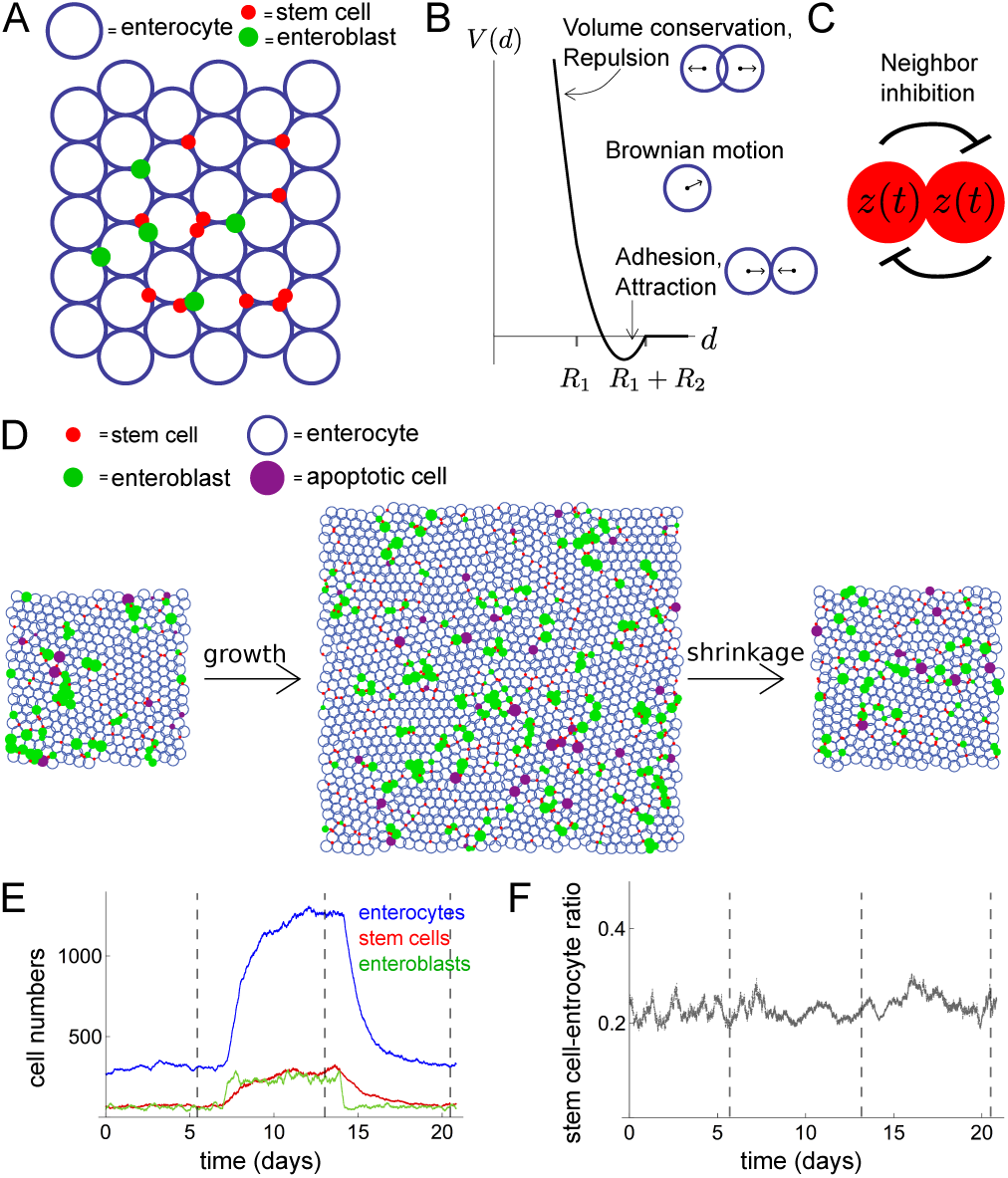
Two-dimensional model accounts for spatial tissue dynamics. Midgut cells are represented by physically interacting, soft pseudo-spheres that signal to each other via Delta-Notch interactions. (A) Schematic: stem cells, enteroblasts, and enterocytes in a 2D domain. (B) Attractive and repulsive forces from neighbor cells and stochastic motility are taken into account. Trapping potential *V* (*d*) from neighbor interactions with preferred distance *d*_0_ between cells of radii *R*_1_ and *R*_2_. (C) Schematic of Delta-Notch lateral inhibition from protein dynamics *z*(*t*). (D) Frames from 2D simulation of feed-fast cycle: *E*_in_ is increased 4-fold then decreased to original value. (E-F) Cell numbers (E) and stem cell-enterocyte ratio (F) as a functions of time for the simulation in (D) with times of frames indicated (vertical dotted lines).

#### Physical cell-cell forces

Assuming that the system is highly damped, we set the inertial term to 0 so that external forces are balanced by viscous drag: *d***x***/dt* = *η***f** where the quantity **f** is the external force acting on a cell, the quantity **x** is the position of the cell center, and *η* is the mobility:

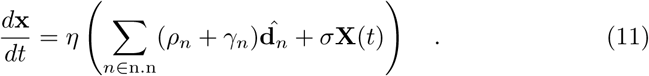

Here, the sum on *n ∈* n.n indicates summing over nearest neighboring cells; if the cell centered at **x** has radius *R*_1_ and another cell has radius *R*_2_, the two cells are nearest neighbors if their cell centers are separated by less than *R*_1_ + *R*_2_. The quantities *ρ*_*n*_ and *γ*_*n*_ are magnitudes of repulsion and adhesion forces from neighbor cells, known to be important in epithelial cells, and **d**_*n*_ is a unit vector towards the neighbor cell. The self-generated random force is provided by **X**(*t*) which has components sampled from a normal distribution *N* (0, 1); here *σ* indicates an amplitude factor for **X**. We refer to *σ***X** as an intrinsic stochastic motility. Cell motility has recently been shown to occur in enteroblasts [53], and labeled clones have been shown to split [15, 16]. The latter may be due to stochastic motility or random cell shuffling, both of which are captured by the term *σ***X**. Forces are indicated in Figure 3B. Details are given in the Supplement.

#### Delta-Notch signaling

In the midgut, asymmetric fate outcomes arise through activation of Notch receptor on the surface of one stem cell by Delta ligand on an adjacent stem cell [33–36, 52, 54]. Activation of Notch marks a cell’s commitment to differentiate [52], and is the defining feature of enteroblast identity. Many mathematical models of Delta-Notch interactions separately describe Delta and Notch populations [54–57], or distinguish between membrane-bound and intracellular Notch [56, 57]. To model lateral inhibition signaling, we employed a fictitious protein with values *z*(*t*) indicating the “stemness” of a given cell. Delta-Notch interactions between neighboring cells are modeled with the dynamics of *z* through a delay differential equation. Delta-Notch signaling have been modeled to contain time delays to account for signal transduction [58], and a delayed model of lateral inhibition has been shown to reduce errors in patterning [59].

For each cell (stem cells and enteroblasts) expressing *z*, we assume that *z* is produced within the cell in a non-linear fashion with saturating effects at level *p*, and *z* decays exponentially with rate *β*. To take into account the effects of nearest neighboring cells, we assume that neighbor cells’ protein levels *z*_*n*_ decreases *z* production in a given cell with time delay *t*_*n*_, taking into account protein transport times, see Figure 3C. Choosing a sigmoidal Hill function of order *m >* 1 for the production term, we have for the dynamics of non-dimensionalized *z*:

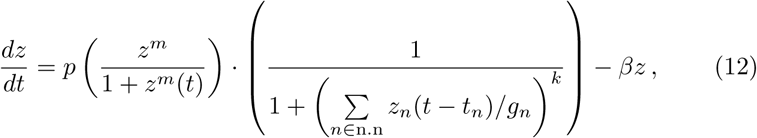

where *g*_*n*_ denotes the switch point for neighbor interactions, and *m* and *k* are Hill exponents. Given that *m >* 1, Equation 12 gives bistable steady states of *z* with stable fixed points at *z*^*\**^ = 0 and *z*^*\**^ *≠* 0 and an unstable fixed point in between that we denote by 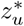. The non-zero stable fixed point *z*^*\**^ *≠* 0 is interpreted as the ‘stem cell’ state, and the trivial stable fixed point *z*^*\**^ = 0 is interpreted as the ‘enteroblast’ state. A stem cell (that has *z >* 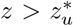) is defined to have permanently “committed” to terminal fate and become an enteroblast when its *z* value falls below the value of the unstable fixed point (*z* becomes 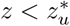). Inhibitory signaling from nearest neighbors cause a stem cell’s *z* value to be suppressed below 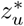, causing commitment. Therefore, in the 2D model, commitment kinetics and commitment rate are determined by dynamics that drive *z* below a threshold value. An enteroblast has a small value of *z* compared to stem cells and therefore signals weakly compared to stem cells; it undergoes growth in size to eventually reach the size of an enterocyte; at this point, the enteroblast is converted into an enterocyte, and its *z* value is set identically to 0. Note that, commitment is still an uncorrelated Poisson process, and therefore, the fraction of symmetric fate outcomes in steady state is still *P* (sym) = 1*/*4. See Supplement for details of implementation of the 2D model.

### Results of 2-dimensional model

Simulations with this 2D model successfully recapitulate essential features of the *in vivo* fly midgut with respect to growth, shrinking, scaling with food, and the presence of stem cells in every part of the epithelium (Figure 3D-F and Movie S1). Additionally, cell populations behave qualitatively similar to the non-spatial *b* = *B*_0_*s/U* model in that they are non-oscillatory (similar to Figure 2D instead of to Figure 2C).

Note that unlike the non-spatial model, the commitment rate *b* for the 2D model is not defined as a function of cell populations. Rather, for each individual cell, commitment rate arises from the duration and strength of contact-based cellcell signaling. This dependence implies that global quantities such as stem cell ratios are determined by local dynamics.

#### Stem cell ratio depends on cells’ physical properties

With the 2D model, we can explicitly examine the effects of local dynamics on population-level quantities by changing physical parameters. Specifically, the forces “adhesion” *γ* and “motility” *σ* in Equation 11 determine how fast stem cells move in the tissue. In particular, stochastic motility specifies diffusive motion absent of other interactions. Additionally, these parameters affect the amount of time stem cells are in signaling contact with other, ultimately affecting the stem cell ratio. Specifically, we found that the stem cell ratio is sensitive to adhesion values, see Figure 4A. Here the motilities *σ*_sc,eb_ for stem cells and enteroblasts and *σ*_EC_ for enterocytes were kept constant while adhesions were varied. Forces are expressed in units of 𝓁 _0_*/*(min*η*), where 𝓁 _0_ is the enterocyte diameter without adhesive or repulsive interactions, and *η* is the inverse viscosity. Note that we use minutes as the unit of time in order to employ the relevant values of *a*_*m*_ and Λ_*m*_ that were deduced from non-spatial model (Figure 2B) to compare to physiological timescales found in experiments.

**Figure 4:**
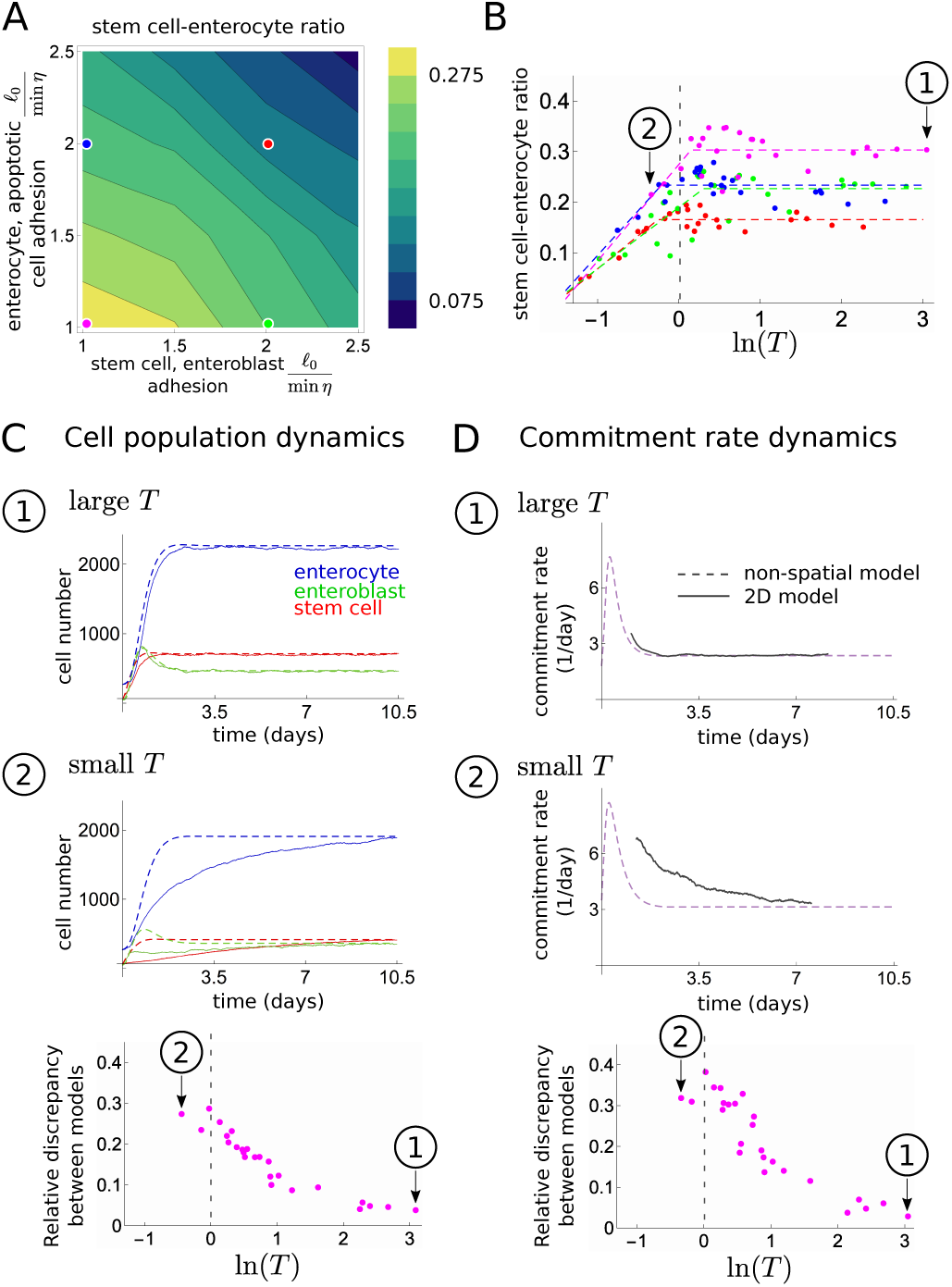
Local physical interactions define a critical ‘territory’ above which stem cells can ‘sense’ their density. (A) Stem cell-enterocyte ratio as a function of adhesion between stem cells and enteroblasts (*x*-axis) and adhesion between enterocytes and apoptotic cells (*y*-axis) for motility *σ* = 0.2𝓁_0_*/*min*η* for all cells. Colored dots indicate adhesion values used in panel (B). (B) Stem cell-enterocyte ratio as a function of natural log of stem cell territory ln(*T*) for various adhesion values. Onset of saturating values for stem cell ratios (dotted line *T ≈* 1) corresponds to lower threshold of *T* that enables stem cell scaling. (C-D) Comparison of 2D (solid lines, average of 6 replicate simulations) and nonspatial *b* = *B*_0_*s/U* model (dashed lines) using time courses of cell population dynamics (C), and commitment rate dynamics (D). Plots correspond to magenta parameter set in (B). Circled numbers indicate large vs. small *T* simulations: large *T* simulations (1) fit well to *b* = *B*_0_*s/U* model while small *T* simulations (2) do not. Calculations of relative discrepancy between models are given in the Supplement.

#### ‘Stem cell territory’ determines whether system scales

In these simulations, we noted that values for adhesion and motility influenced the physical space that a stem cell explores during its lifetime. We denote this space as a ‘stem cell territory’, *T*. To gain insight into the relationship between territories and stem cell interactions, we define *T* as:

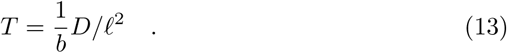

Here, 1*/b* is the average lifetime of a stem cell, i.e., the time between the birth of a stem cell and its commitment to terminal fate; the quantity *D* is a diffusion constant fitted to simulated stem cell tracks (see Supplement); and *f* is the average enterocyte diameter. Hence, *T* is the dimensionless area (number of enterocyte areas) over which a stem cell may influence or respond to other stem cells. Note that Equation 13 can be generalized for non-diffusive motion.

Investigating resulting stem cell ratios as a function of *T* (Figure 4B), we find a transition from increasing stem cell ratios at lower *T* towards saturation for larger *T*. Here, the quantity *T* was varied via adhesion *γ* and stochastic motility *σ*: as expected, larger *σ* and lower *γ* lead to larger territories *T*. Note that this transition occurs around *T ≈* 1, i.e. when stem cells start to explore a territory larger than the area of a single enterocyte. For *T >* 1, different stem cells can come in contact with each other via overlapping territories. The saturation for large *T* also indicates that additional interactions do not change global behavior. Moreover, the resulting stem cell ratio for large *T* depends on the intrinsic parameters (adhesion *γ*); this dependence suggests that the tissue can flexibly regulate its stem cell ratio.

It is insightful to compare the 2D simulations to the non-spatial model Figure 4 and Equation 10. We find that for large stem cell territories *T*, the population dynamics of the two models agree (simulation (1) in Figure 4C-D). This suggests that when *T* is large, stem cells can ‘sense’ their density within the tissue, and hence their commitment rate can contain density feedback through *s/U*, as in the non-spatial model. When *T* is small, the two models differ (simulation (2) in Figure 4C-D) since the non-spatial model cannot account for the effect, in 2D, that stem cells with small territories do not obtain density knowledge. Additionally, we note that in the latter case, when *T* is small, the approximate time to reach new homeostatic cell numbers during resizing is longer than observed physiologically (approximately 7 days instead of 3.5 days [15]); this supports the notion that stem cells in the biological system may display motility to increase their territories.

Finally, we explored how stem cell territory *T* relates to stem-cell scaling. We performed numerical experiments for different values of food input *E*_in_ for various *T* (see the Supplement and Figure S4). We found that the regime for which stem cell scaling is enabled approximately coincides with the regime for which stem cell ratios saturate: at *T ≥* 1 (dotted vertical line in Figure 4B). Therefore, when stem cells explore a large enough area to escape the inhibition signaling of its immediate sibling stem cell, stem cell scaling is enabled.

## Discussion

Proportional scaling of stem cells to total cells during adaptive growth ensures that the organ has enough stem cells to support its new size after growth is complete. We have shown that the basic features of adaptive organ resizing can be captured by simple mathematical descriptions of the Drosophila midgut in which rates of stem cell division and enterocyte death depend on nutrient density. Importantly, we find that stem cells must commit to a terminal fate at a non-constant rate, and that a rate tuned to stem cell proportion reproduces the *in vivo* kinetics of division and growth.

What biological mechanisms might enable cells to ‘monitor’ stem cell proportions and tune their commitment rates appropriately? The models presented here show that a mechanism involving stochastic motility is compatible with *in vivo* measurements. Motility permits stem cells to explore a local tissue area and engage in signaling interactions with other, potentially non-sibling, stem cells. We define this local area as the stem cell’s ‘territory’ and explore its parameter space by varying cell motility and adhesion.

Intriguingly, proportion-dependent commitment and scaling occur only above a threshold territory size. This threshold size can be understood by its impact on commitment rate (i.e., rate of Notch activation), which occurs through cell-cell signaling (Notch-Delta interactions) between pairs of stem cells. Essentially, there are two scenarios: (1) When stem cells are constrained within small territories, two newborn sibling cells contact each other frequently, and their Notch-Delta interactions increase to the point of a near-certain asymmetric outcome. In this case, the total number of stem cells in the tissue remains nearly constant during growth, while the number of enterocytes increases, and scaling does not occur (Figure 5A,C). (2) When stem cells range in large territories, sibling cells often separate fast enough without Notch-Delta interaction having a differentiating effect, which promotes symmetric fate outcomes, i.e., the total number of stem cells in the tissue can increase. Importantly, the stem cell ratio does not increase beyond a certain value since pairs of non-sibling stem cells come into contact sufficiently frequently for Notch-Delta interactions to induce differentiation. In this case, both stem cells and enterocytes increase in proportion, and scaling does occur (Figure 5B,D).

**Figure 5:**
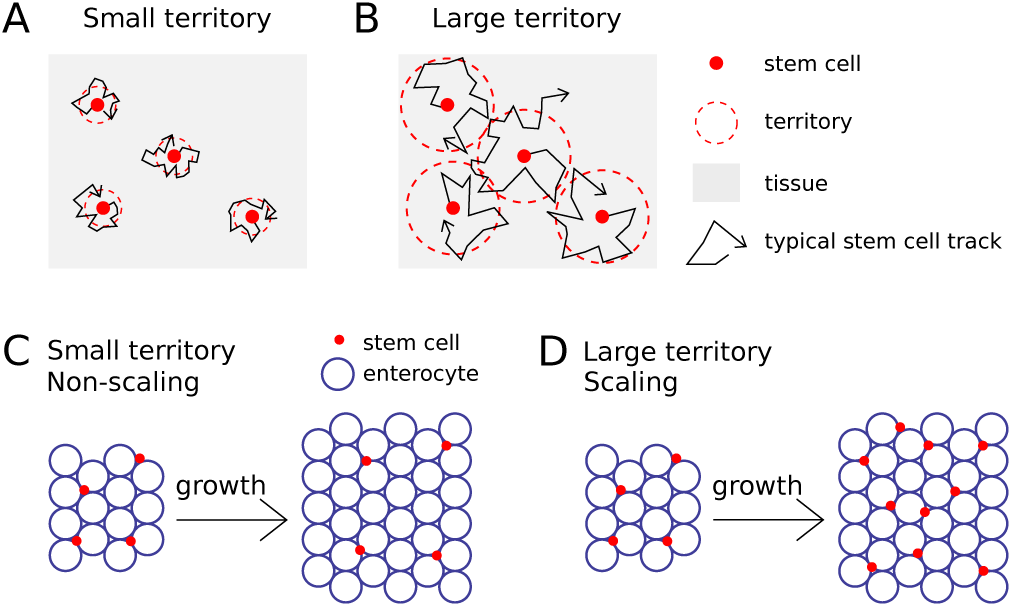
Scaling occurs when stem cell territories are above a critical threshold. (A-B) Cartoons of small (A) versus large (B) stem cell territory (red dotted line) as defined by spatial range of physical cell motion (black line). (C) For territories below a threshold size, there is no stem cell scaling. (D) For territories larger than a threshold size, there is stem cell scaling.

We make the following suggestions for future experimental tests. The core importance of a tunable commitment rate in these models contrasts with the scant empirical knowledge of commitment rates *in vivo*. Experimental measurements of commitment rate are currently impractical in many systems. However, the availability of fluorescent Notch reporters [60, 61] may provide a means to measure kinetics of fate commitment in systems such as the midgut in which Notch activation is the committing step. For the midgut in particular, the models here generate specific, testable predictions: (1) If newborn stem cells inherit unequal levels of the Delta ligand or Notch receptor, then because of the delayed nature of Equation 12, commitment should occur more rapidly in sibling pairs than in pairs of stem cells that come into contact by chance. (2) The stem cell population should undershoot when a tissue undergoes shrinkage (Figure 2D). (3) Stem cell scaling should be disrupted by experimental perturbation of stochastic motility, adhesive force, or Delta-Notch signaling (Figure 4B). (4) The territory size of individual stem cells should be larger than one enterocyte area. In particular, for the last point (4), methods such as cell tracking *in vivo* to measure diffusion coefficients of cell motion, or clone-induction methods such as twin-spot mosaic analysis [15] to measure spatial dispersal of cells from a common division, can provide measurements and estimations of the stem cell territory in the midgut.

In summary, we have developed mathematical descriptions of a stem cell-based organ that undergoes adaptive resizing in response to external input. To realistically describe an *in vivo* system, the Drosophila midgut, we found that the stem cell commitment rate should depend on organ-wide stem cell proportion. To elucidate this dependence, we suggested local, spatially-motivated, cell-level mechanisms such as cell motility, adhesion, and signaling, by which stem cells can detect their density and therefore tune their commitment rate. Importantly, these models naturally give rise to stem cell scaling, and we identify physical regimes in which scaling occurs.

## Author Contributions

X.D, L.E.O, and I.R.K designed research, performed research, contributed analytic tools, analyzed data, and wrote the manuscript.

## Acknowledgements

We thank S. Streichan, A. Lam, and T. Petty for valuable comments on the manuscript, and members of I.R.K. and L.E.O. labs for useful discussions.

## References

[1] T. Piersma and Å. Lindström, “Rapid reversible changes in organ size as a component of adaptive behaviour,” Trends in Ecology & Evolution 12 (1997) 134–138.

[2] H. S. Aldewachi, N. A. Wright, D. R. Appleton, and A. J. Watson, “The effect of starvation and refeeding on cell population kinetics in the rat small bowel mucosa,” Journal of Anatomy 119 (1975) 105–121.

[3] G. G. Altmann, “Influence of starvation and refeeding on mucosal size and epithelial renewal in the rat small intestine,” American Journal of Anatomy 133 (1972) 391–400.

[4] M. J. Koury, “Erythropoietin: the story of hypoxia and a finely regulated hematopoietic hormone,” Experimental hematology 33 (2005), no. 11 1263–1270.

[5] H. O. Brown, M. L. Levine, and M. Lipkin, “Inhibition of intestinal epithelial cell renewal and migration induced by starvation,” American Journal of Physiology 205 (1963) 868–872.

[6] S. Dunel-Erb, C. Chevalier, P. Laurent, A. Bach, F. Decrock, and Y. Le Maho, “Restoration of the jejunal mucosa in rats refed after prolonged fasting,” Comparative Biochemistry and Physiology Part A: Molecular & Integrative Physiology 129 (2001), no. 4 933–947.

[7] L. A. Meyers and J. J. Bull, “Fighting change with change: adaptive variation in an uncertain world,” Trends in Ecology & Evolution 17 (2002) 551–557.

[8] H. V. Carey, “Seasonal changes in mucosal structure and function in ground squirrel intestine,” American Journal of Physiology-Regulatory, Integrative and Comparative Physiology 259 (1990), no. 2 R385–R392.

[9] S. M. Secor and J. Diamond, “A vertebrate model of extreme physiological regulation,” Nature 395 (1998), no. 6703 659–662.

[10] N. A. Dumont, C. F. Bentzinger, M.-C. Sincennes, and M. A. Rudnicki, “Satellite cells and skeletal muscle regeneration,” Comprehensive Physiology (2015).

[11] F. Ambrosio, F. Kadi, J. Lexell, G. K. Fitzgerald, M. L. Boninger, and J. Huard, “The effect of muscle loading on skeletal muscle regenerative potential: an update of current research findings relating to aging and neuromuscular pathology,” American journal of physical medicine & rehabilitation/Association of Academic Physiatrists 88 (2009), no. 2 145.

[12] S. Schiaffino, S. P. Bormioli, and M. Aloisi, “Cell proliferation in rat skeletal muscle during early stages of compensatory hypertrophy,” Virchows Archiv B 11 (1972), no. 1 268–273.

[13] J. E. Visvader, “Keeping abreast of the mammary epithelial hierarchy and breast tumorigenesis,” Genes & development 23 (2009), no. 22 2563–2577.

[14] A. C. Rios, N. Y. Fu, G. J. Lindeman, and J. E. Visvader, “In situ identification of bipotent stem cells in the mammary gland,” Nature 506 (2014), no. 7488 322–327.

[15] L. E. O’Brien, S. S. Soliman, X. Li, and D. Bilder, “Altered modes of stem cell division drive adaptive intestinal growth,” Cell 147 (2011) 603–614.

[16] J. de Navascués, C. N. Perdigoto, Y. Bian, M. H. Schneider, A. J. Bardin, Martínez-Arias, and B. D. Simons, “*Drosophila* midgut homeostasis involves neutral competition between symmetrically dividing intestinal stem cells,” The EMBO Journal 31 (2012), no. 11 2473–2485.

[17] S. Itzkovitz, I. C. Blat, T. Jacks, H. Clevers, and A. van Oudenaarden, “Optimality in the Development of Intestinal Crypts,” Cell 148 (2012) 608–619.

[18] E. Hannezo, J. Prost, and J.-F. Joanny, “Growth, homeostatic regulation and stem cell dynamics in tissues,” Journal of the Royal Society: Interface 11 (2014) 1–10.

[19] S. Rulands and B. D. Simons, “Tracing cellular dynamics in tissue development, maintenance and disease,” Current Opinion in Cell Biology 43 (2016) 38–45.

[20] T. Krieger and B. D. Simons, “Dynamic stem cell heterogeneity,” Development 142 (2015), no. 8 1396–1406.

[21] B. D. Simons and H. Clevers, “Strategies for Homeostatic Stem Cell Self-Renewal in Adult Tissues,” Cell 145 (2011) 851–862.

[22] H. J. Snippert, L. G. van der Flier, T. Sato, J. H. van Es, M. van den Born, C. Kroon-Veenboer, N. Barker, A. M. Klein, J. van Rheenen, B. D. Simons, and H. Clevers, “Intestinal crypt homeostasis results from neutral competition between symmetrically dividing Lgr5 stem cells,” Cell 143 (2010) 134–144.

[23] A. M. Klein and B. D. Simons, “Universal patterns of stem cell fate in cycling adult tissues,” Development 138 (2011) 3103–3111.

[24] A. M. Klein, D. P. Doupé, P. H. Jones, and B. D. Simons, “Kinetics of cell division in epidermal maintenance,” Physical Review E 76 (2007) 021910.

[25] A. M. Klein, D. P. Doupé, P. H. Jones, and B. D. Simons, “Mechanism of murine epidermal maintenance: Cell division and the voter model,” Physical Review E 77 (2008) 031907.

[26] I. M. Van Leeuwen, G. Mirams, A. Walter, A. Fletcher, P. Murray, J. Osborne, S. Varma, S. Young, J. Cooper, B. Doyle, *et. al*., “An integrative computational model for intestinal tissue renewal,” Cell proliferation 42 (2009), no. 5 617–636.

[27] M. D. Johnston, C. M. Edwards, W. F. Bodmer, P. K. Maini, and S. J. Chapman, “Mathematical modeling of cell population dynamics in the colonic crypt and in colorectal cancer,” Proceedings of the National Academy of Sciences 104 (2007), no. 10 4008–4013.

[28] S. Kunche, H. Yan, A. L. Calof, J. S. Lowengrub, and A. D. Lander, “Feedback, Lineages and Self-Organizing Morphogenesis,” PLoS Comput Biol 12 (2016), no. 3 e1004814.

[29] P. Rué, Y. H. Kim, H. L. Larsen, A. Grapin-Botton, and A. M. Arias, “A framework for the analysis of symmetric and asymmetric divisions in developmental process,” BioArxiv (2015) 1.

[30] M. P. Alcolea, P. Greulich, A. Wabik, J. Frede, B. D. Simons, and P. H. Jones, “Differentiation imbalance in single oesophageal progenitor cells causes clonal immortalization and field change,” Nature Cell Biology 16 (2014) 615–622.

[31] G. Kolahgar, S. J. Suijkerbuijk, I. Kucinski, E. Z. Poirier, S. Mansour, B. D. Simons, and E. Piddini, “Cell Competition Modifies Adult Stem Cell and Tissue Population Dynamics in a JAK-STAT-Dependent Manner,” Developmental cell 34 (2015), no. 3 297–309.

[32] T. Thalheim, P. Buske, J. Przybilla, K. Rother, M. Loeffler, and J. Galle, “Stem cell competition in the gut: insights from multi-scale computational modelling,” Journal of The Royal Society Interface 13 (2016), no. 121 20160218.

[33] B. Ohlstein and A. Spradling, “The adult *Drosophila* posterior midgut is maintained by pluripotent stem cells,” Nature Letters 439 (2006) 470–474.

[34] C. A. Micchelli and N. Perrimon, “Evidence that stem cells reside in the adult *Drosophila* midgut epithelium,” Nature 439 (2006) 475–479.

[35] B. Ohlstein and A. Spradling, “Multipotent *Drosophila* Intestinal Stem Cells Specify Daughter Cell Fates by Differential Notch Signaling,” Science 315 (2007) 988–992.

[36] A. J. Bardin, C. N. Perdigoto, T. D. Southall, A. H. Brand, and F. Schweisguth, “Transcriptional control of stem cell maintenance in the Drosophila intestine,” Development 137 (2010), no. 5 705–714.

[37] R. Farhadifar, J.-C. Röper, B. Aigouy, S. Eaton, and F. Jülicher, “The influence of cell mechanics, cell-cell interactions, and proliferation on epithelial packing,” Current Biology 17 (2007), no. 24 2095–2104.

[38] T. Nagai, K. Kawasaki, and K. Nakamura, “Vertex dynamics of two-dimensional cellular patterns,” Journal of the Physical Society of Japan 57 (1988), no. 7 2211–2224.

[39] H. Honda, Y. Ofita, S. Higuchi, and K. Kani, “Cell movements in a living mammalian tissue: long-term observation of individual cells in wounded corneal endothelia of cats,” Journal of Morphology 174 (1982) 25–39.

[40] K. Kawasaki, T. Nagai, and K. Nakashima, “Vertex models for two-dimensional grain growth,” Philosophical Magazine Part B 60 (1989), no. 3 399–421.

[41] M. J. Simpson, A. Merrifield, K. A. Landman, and B. D. Hughes, “Simulating invasion with cellular automata: connecting cell-scale and population-scale properties,” Physical Review E 76 (2007), no. 2 021918.

[42] A. R. Kansal, S. Torquato, G. Harsh, E. Chiocca, and T. Deisboeck, “Simulated brain tumor growth dynamics using a three-dimensional cellular automaton,” Journal of theoretical biology 203 (2000), no. 4 367–382.

[43] Y. Lee, S. Kouvroukoglou, L. V. McIntire, and K. Zygourakis, “A cellular automaton model for the proliferation of migrating contact-inhibited cells,” Biophysical journal 69 (1995), no. 4 1284–1298.

[44] F. Graner and J. A. Glazier, “Simulation of biological cell sorting using a two-dimensional extended Potts model,” Physical review letters 69 (1992), no. 13 2013.

[45] J. M. Osborne, “Multiscale model of colorectal cancer using the cellular potts framework,” Cancer informatics 14 (2015), no. Suppl 4 83.

[46] P. Buske, J. Galle, N. Barker, G. Aust, H. Clevers, and M. Loeffler, “A comprehensive model of the spatio-temporal stem cell and tissue organisation in the intestinal crypt,” PLoS Comput Biol 7 (2011), no. 1 e1001045.

[47] J. M. Osborne, A. G. Fletcher, J. M. Pitt-Francis, P. K. Maini, and D. J. Gavaghan, “Comparing individual-based approaches to modelling the self-organization of multicellular tissues,” PLOS Computational Biology 13 (2017), no. 2 e1005387.

[48] M. Basan, J. Elgeti, E. Hannezo, W.-J. Rappel, and H. Levine, “Alignment of cellular motility forces with tissue flow as a mechanism for efficient wound healing,” Proceedings of the National Academy of Sciences 110 (2013), no. 7 2452–2459.

[49] D. Drasdo, R. Kree, and J. S. McCaskill, “Monte Carlo approach to tissue-cell populations,” Physical Review E 52 (1995), no. 6 6635–6657.

[50] D. Drasdo and S. Höhme, “A single-cell-based model of tumor growth in vitro: monolayers and spheroids,” Physical Biology 2 (2005) 133–147.

[51] E.-M. Schötz, M. Lanio, J. A. Talbot, and M. L. Manning, “Glassy dynamics in three-dimensional embryonic tissues,” Journal of Royal Society Interface 10 (2013) 1–11.

[52] C. N. Perdigoto, F. Schweisguth, and A. J. Bardin, “Distinct levels of Notch activity for commitment and terminal differentiation of stem cells in the adult fly intestine,” Development 138 (2011) 4585–4595.

[53] Z. A. Antonello, T. Reiff, E. Ballesta-Illan, and M. Dominguez, “Robust intestinal homeostasis relies on cellular plasticity in enteroblasts mediated by miR-8– Escargot switch,” The EMBO Journal May (2015) 1.

[54] N. Guisoni, R. Martinez-Corral, J. Garcia-Ojalvo, and J. de Navascues, “Diversity of fate outcomes in cell pairs under lateral inhibition,” Development 144 (2017), no. 7 1177–1186.

[55] J. R. Collier, N. A. Monk, P. K. Maini, and J. H. Lewis, “Pattern Formation by Lateral Inhibition with Feedback: a Mathematical Model of DeltaNotch Intercellular Signalling,” Journal of Theoretical Biology 183 (1996) 429–446.

[56] D. Sprinzak, A. Lakhanpal, L. LeBon, L. A. Santat, M. E. Fontes, G. A. Anderson, J. Garcia-Ojalvo, and M. B. Elowitz, “Cis-interactions between Notch and Delta generate mutually exclusive signalling states,” Nature Letters 456 (2010) 86–91.

[57] D. Sprinzak, A. Lakhanpal, L. LeBon, J. Garcia-Ojalvo, and M. B. Elowitz, “Mutual Inactivation of Notch Receptors and Ligands Facilitates Developmental Patterning,” PLoS Computational Biology 7 (2011) e1002069.

[58] O. Barad, D. Rosin, E. Hornstein, and N. Barkai, “Error minimization in lateral inhibition circuits,” Sci Signal 3 (2010), no. 129 ra51.

[59] D. S. Glass, X. Jin, and I. H. Riedel-Kruse, “Signaling delays preclude defects in lateral inhibition patterning,” Physical review letters 116 (2016), no. 12 128102.

[60] S. Barolo, B. Castro, and J. Posakony, “New Drosophila transgenic reporters: insulated P-element vectors expressing fast-maturing RFP.,” BioTechniques 36 (2004), no. 3 436–40.

[61] B. E. Housden, K. Millen, and S. J. Bray, “Drosophila reporter vectors compatible with ΦC31 integrase transgenesis techniques and their use to generate new Notch reporter fly lines,” G3: Genes—Genomes—Genetics 2 (2012), no. 1 79–82.

